# Dissociation of novel open loop from ventral putamen to motor areas from classic closed loop in humans II: task-based function

**DOI:** 10.1101/2024.06.18.599622

**Authors:** Neil M. Dundon, Elizabeth Rizor, Joanne Stasiak, Jingyi Wang, Kiana Sabugo, Christina Villaneuva, Parker Barandon, Andreea C. Bostan, Regina C. Lapate, Scott T. Grafton

## Abstract

Humans ubiquitously increase the speed of their movements when motivated by incentives (i.e., capturing reward or avoiding loss). The complex interplay between incentivization and motor output is pertinent for unpacking the functional profiles of different circuits that link the basal ganglia with motor cortical areas. Here, we analyzed the functional profile of nodes forming two circuits involving putamen and motor cortical areas: the traditional “closed-loop circuit” (CLC) from sensorimotor dorsal putamen (PUTd) and a putative “open-loop circuit” (OLC) from ventral putamen (PUTv). Establishing differential function between CLC and OLC is particularly relevant for therapeutic approaches to Parkinson’s disease, where OLC function is hypothesized to be relatively spared by the disease process. In a large sample fMRI study, 68 healthy controls executed speeded reaches with a joystick under different levels of incentivization to accurately hit precision targets. We dissociated effects of “incentive per se” (i.e., changes in brain activity when an upcoming movement obtains a reward or avoids a loss) from “RT effects” (i.e., brain activity that directly scales with adjustments to movement initiation time). Incentive per se was observed across sites in both CLC and OLC. However, RT effects were primarily in nodes of the OLC and motor sites, consistent with the hypothesized anatomy and function of OLC. Our findings additionally suggest valence might mediate when incentives recruit OLC to more prominent control of motor behavior.

## Introduction

One of the most important—if not the most important—goal-directed behavior is initiating a physical movement. Traditional neuroanatomical understanding of movement initiation in mammals centers on reciprocal projections between the primary motor cortex (M1) and the basal ganglia. The canonical circuit consists of corticostriatal projections into basal ganglia via dorsal portions of the caudal putamen (sensorimotor putamen; PUTd). The circuit proceeds via PUTd projections to globus pallidus internus (GPi), the primary output nucleus of basal ganglia. GPi then projects back to M1 via projections with ventral sites in the motor thalamus. These cortico-striato-thalamo-cortical projections thus comprise a “closed-loop” circuit (CLC). Its crucial role in facilitating movement initiation is highlighted by the profound deficits (bradykinesia, akinesia, rigidity) observed in Parkinson’s disease (PD)—a disease characterized by targeted depletion of dopaminergic inputs to PUTd in particular.

However, transneuronal-tracer findings suggest the presence of an alternative circuit through basal ganglia that might facilitate movement initiation, without forming a closed loop. First, retrograde transneuronal transport of rabies virus in nonhuman primates has revealed that ventral sites in putamen (PUTv) send projections to M1, even though they do not receive reciprocal innervation from it. These PUTv sites instead receive projections from amygdala and limbic cortical areas (1, 2), suggesting that an “open-loop” circuit (OLC; Figure 1A) might link limbic regions of the putamen to motor areas (3–6). In addition, PUTv shows spared dopaminergic innervation in primate models of PD (7), consistent with observations in human patients where dopamine-agonizing medications can disrupt motor function by way of an “overdose” effect during tasks (sequence learning) that recruit PUTv (8). Another phenomenon occasionally seen in PD patients is “Paradoxical Kinessa” (PK; (9), whereby emotionally arousing circumstances such as car accidents (10) or natural disasters (11, 12) can seemingly drive transient normalization of motor function (13).

**Figure 1.**
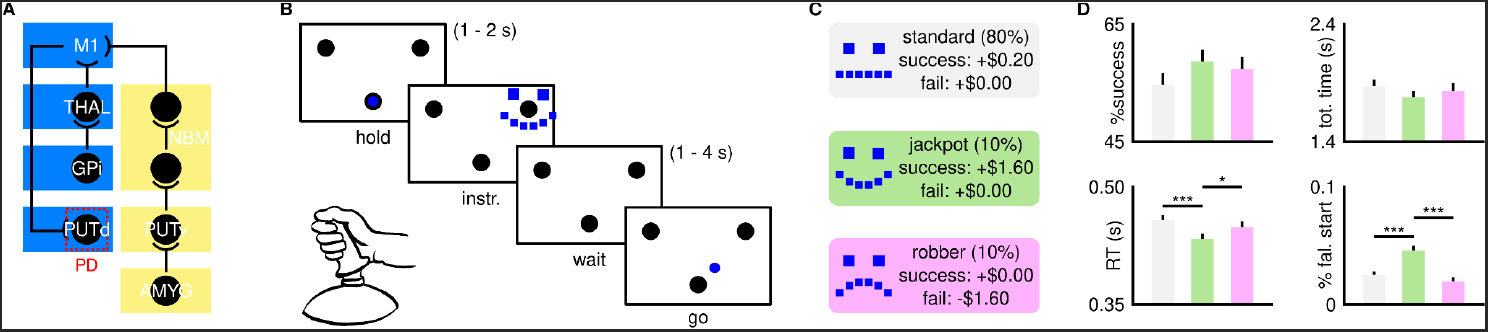
**A** Closed (CLC;blue) and open (OLC;yellow) motor loops. **B** Incentivized reaching task, **C** incentive conditions and **D** behavioral results.

Together, these findings suggest that the putative OLC might exist to afford limbic and cognitive processes an influence over movement initiation in certain contexts. However, it remains unclear whether specific contextual factors lead to stronger associations between movement initiation and nodes of the putative OLC compared to those in the canonical CLC. We accordingly report data from a task-based functional magnetic resonance (fMRI) experiment aimed at functionally dissociating CLC from putative OLC in humans. We present results from a large sample (n=68) of participants performing speeded manual precision reaches under different contexts of incentivization (positive and negative valence). We used state-of-the-art modeling of the accompanying fMRI-derived brain signals (14) that allowed us to separate regions associated with incentive independently of behavior changes (incentive per se) from those associated with the incentive-derived changes in movement initiation time (RT effects). We see that nodes of both CLC and OLC circuits activate more under different contexts of incentive per se, however, RT effects were more prominent in nodes of the putative OLC. We additionally see that RT effects in OLC are more prevalent with incentives of positive rather than negative valence, and associated with motor regions consistent with the proposed structure and function of this circuit.

## Results

68 human participants completed three runs of an incentivized reaching task during fMRI. This sample included 50 females and were an average age of 20.75 (SD=1.86). Each run contained 100 trials in which they completed a precision reach to a cued target with a computer joystick (Figure 1B). To successfully complete a trial, reaches needed to enter and be held inside the target for one second before a deadline. The deadline for reach completion from onset of the go cue (1.87 s) was calibrated from a pilot sample to achieve a 50% success rate across participants. Each trial began with participants holding a screen cursor (yoked to joystick position) at a starting position. An instruction cue then appeared at one of two spatial locations, i.e., the spatial target for that trial. The identity of the instruction cue also communicated the trial’s incentive. On 80% of trials (“standard” trials) participants earned $0.20 if successful, or $0.00 if unsuccessful. The remaining 20% of trials provided “high incentive”. Half of these (i.e., 10% of total) trials were “jackpot” trials, offering high positive valence incentive. On such trials, participants earned $1.60 if successful, or $0.00 if unsuccessful. The remaining trials (10% of total) were “robber” trials, offering high negative valence incentive, where participants avoided a loss of $1.60 if successful. Following a hold period, a go cue informed them to reach for the cued target. Visual feedback informed participants whether or not they had successfully completed the trial prior to the deadline.

Behavioral data verified that high positive valence incentives selectively modulated the expedience of movement initiation. Overall, participants successfully completed 56.8% of trials. However, while success rates were modulated by incentive context (F(2,134)=4.16,p=0.018), no pairwise comparisons of success between contexts were significant after multiple-comparison adjustment (all pairwise p_fdr_>0.087; Figure 1D). Median completion time was 1.66 s for successful trials and 2.40 s for unsuccessful trials, however this metric was not otherwise modulated by incentive context (F(2,134)=0.7049,p=0.496; Figure 1D). Instead, the incentive context modulated median time taken to initiate movement (initiation time—RT; F(2,134)=18.5,p<0.001; Figure 1D), with participants initiating movement on jackpot trials (IT=0.44 s) significantly faster than both standard (0.46 s;p_fdr_<0.001) and robber trials (0.45 s;p_fdr_=0.013). Consistent with high positive valence driving an incentivization to move, over other cognitive components of optimized reaching, participants also committed a “false start” (movement prior to appearance of the go cue) more often on jackpot trials (F(2,134)=25.4;p<0.001; Figure 1D).

### Incentive per se in the brain

We next modeled the accompanying fMRI-derived brain signals. In a first GLM, we sought to identify brain regions where the average voxel activation was greater in high-incentive contexts, while ensuring that any such effects were not confounded by context-general mechanisms that scale activation intensity with initiation time (14). For this, we compared brain activity between jackpot and robber trials versus standard trials, separately for the instruction phase and go phase of trials, i.e., four contrasts (jackpot_inst_>standard_inst_; robber_inst_>standard_inst_; jackpot_go_>standard_go_; robber_go_>standard_go_). A separate contrast using a duration-scaled (by RT) regressor event-locked to the go cue (RT_go_>baseline) then accounted for scaled (i.e., behaviorally relevant) activations across all contexts (Figure 2A). We tested for credible nonzero average voxel activation in all ROIs (Figure 2B) for each contrast using the same hierarchical Bayesian mixture model (Student’s T framework) to control for Type I error and ensure reliable results. We report here on functionally relevant left-lateralized sites, i.e., contralateral to the limb performing reaches.

**Figure 2.**
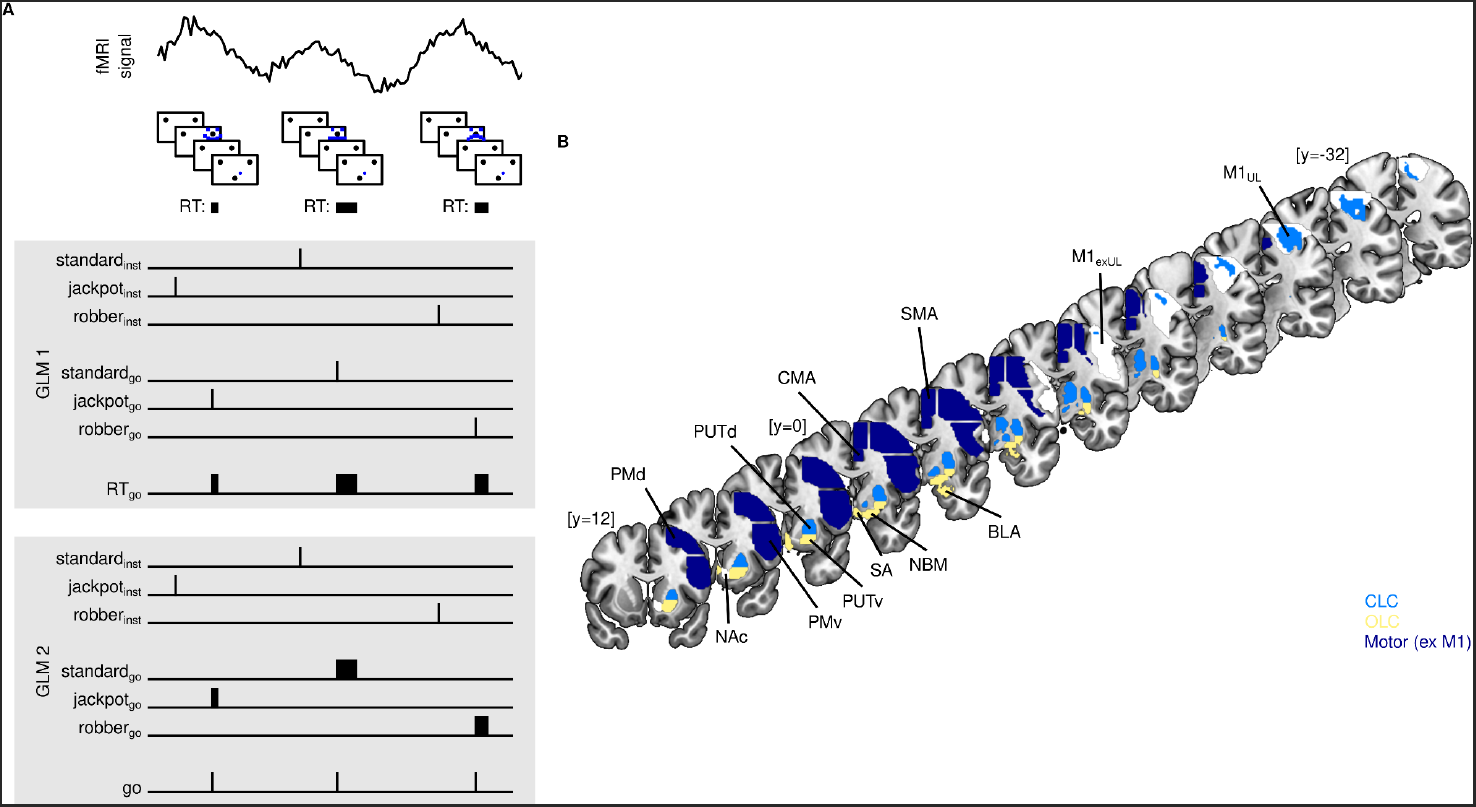
**A** fMRI GLM frameworks and **B** ROI masks (M1exUL shaded while to distinguish from M1_UL_; NAc shaded white as not hypothesized to be in either circuit)

We observed effects of incentive per se across sites of OLC, CLC and motor areas, but at different phases of trials depending on the valence of incentive. On jackpot trials (high positive valence incentive), average voxel activation credibly increased (relative to standard trials) in multiple sites, but solely following instruction cues (jackpot_inst_>standard_inst_). This includes OLC sites in basal forebrain (nucleus basalis of Meynert—NBM, septal region—SA), amygdala (basolateral nucleus—BLA) and PUTv (Figure 3A). From our target OLC sites, only amygdala (central nucleus—CeA) did not show this effect. We also observed this effect of jackpot incentive per se at instruction in CLC sites—in globus pallidus internus (GPi) and ventrolateral thalamus (VL; Figure 3a). From our target CLC sites, only PUTd did not show the effect. We also observed this effect of jackpot incentive per se at instruction in each of our target motor areas—in primary motor cortex (upper-limb region—M1_UL_), primary motor cortex (excluding upper-limb region—M1_exUL_), premotor cortex (dorsal—PMd, ventral—PMv), supplementary motor area (SMA), and cingulate motor area (CMA; Figure 3a). We additionally saw this effect in the nucleus accumbens (NAc). In contrast, on robber trials (high negative valence), at instruction, average voxel activation did not credibly increase (relative to standard trials) in any of our target sites (whether in OLC, CLC or a motor area). In sum, as participants realized that an ensuing movement would yield a high reward (but not avoid a large loss), we observed a broad increase in voxel activations across functionally relevant sites of OLC, CLC and motor areas.

**Figure 3.**
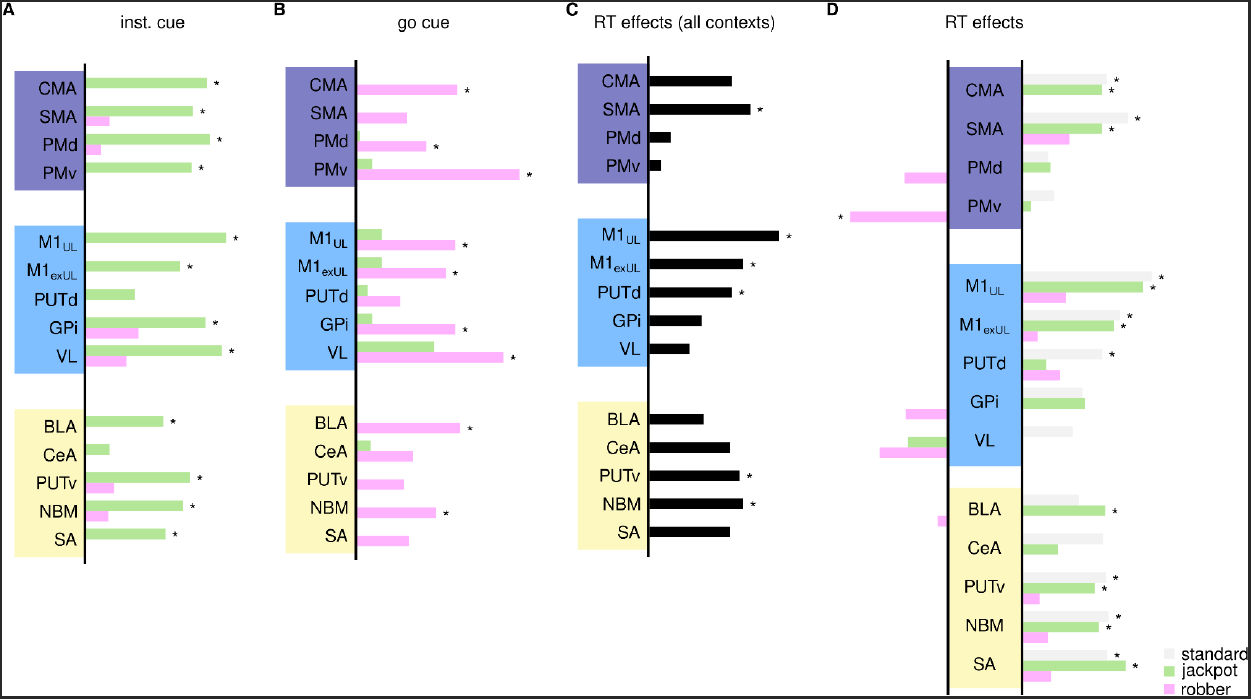
**A-B** Neural activations for incentive per se at instruction (**A**) and go **(B)** phase of trials. **C-D** Neural activations for RT effects collapsed across trials **(C)** and separated by context (**D**). * credibly non-zero activation.

At the go cue, we observed a reversal whereby brain regions activated to negative valence instead. i.e., as participants initiated their goal-directed movement, voxel activations did not credibly increase on jackpot trials in any sites. Instead, at this go phase of trials, activations credibly increased on robber trials in multiple sites. This included OLC sites—NBM and BLA. From target OLC sites, SA, CeA and PUTv did not show this effect (Figure 3B). We also observed this effect of robber incentive per se at go cue in CLC sites—GPi, STN and VL (Figure 3B). From our target CLC sites, only PUTd did not show the effect. We also observed this effect of robber incentive per se at go cue in each of our target motor areas except SMA, that is, we observed the effect in M1_UL_, M1_exUL_, PMd, PMv and CMA (Figure 3B). In sum, as participants initiated a movement to avoid a large loss (but not yield a high reward), we observed a broad increase in the average voxel activation across functionally relevant sites of OLC, CLC and the motor areas.

Merging our results so far regarding incentive per se, we observed an apparent cross-over interaction involving trial phase and valence in a large volume of our tested sites. That is, for multiple sites, activations increased only at the moment of instruction on jackpot trials, and only at the moment of movement initiation (go cue) on robber trials. The specific OLC sites that showed this cross-over interaction were NBM and BLA. Specific CLC sites that showed this cross-over interaction were GPi, STN and VL. Specific motor sites that showed this cross-over interaction were M1_UL_, M1_exUL_, PMd, PMv and CMA. The cross-over effect was also observed in NAc. Notably, a subset of our target sites did not show the cross-over effect. This subset instead showed responsivity which was more reward-specific, with credibly heightened activations only on jackpot trials at the moment of instruction. This subset included two sites in CLC—SA, PUTv, and a single motor area—SMA. In sum, our results so far suggest that in addition to sites showing temporal- and valence-flexible responsivity (i.e., the cross-over interaction effect above), a separate network of OLC and motor areas might respond specifically to reward incentive at the moment participants learn the incentive context.

The above effects (incentive per se) describe greater brain activations for high incentive at different phases of trials, but they do not reveal neural associations with behavioral output. To this latter aim, the RT-scaled regressor included in the same GLM that directly tested for brain activity scaling with adjustments to initiation speed also showed credible associations with a number of sites “RT effects” (Figure 3C). This included OLC sites (NBM, PUTv) and CLC sites (PUTd). We additionally saw RT effects in motor areas (M1_UL_, M1_exUL_ and SMA) and in NAc. All activations were in a direction whereby faster movement initiation was related to greater activation intensity across regions’ voxels. This initial profile of RT effects suggests that alongside brain regions that increase activations at various phases of high incentivization, a number of brain regions also scale their activations with the expedience of movement initiation. In the following section, we report from a second model that parsed RT effects across the different incentive contexts.

### RT effects separated by incentive context

We modeled the brain signals with a separate GLM (Figure 2A) to identify context-specific RT effects, that is, brain regions that were associated with movement initiation under specific incentives. We preserved regressors from the first GLM, allowing contrasts of incentive per se at the instruction phase of trials (i.e., jackpot_inst_>standard_inst_; robber_inst_>standard_inst_). At the go phase of trials, we then inverted the logic of the first GLM, that is, testing for separate RT effects for the three incentive contexts (standard_RTgo_>baseline; jackpot_RTgo_>baseline; robber_RTgo_>baseline) alongside a single stick function to account for go-cue related activations independent of behavior and regardless of context (Go > baseline). We again used the hierarchical Bayesian model testing all contrasts and regions together.

Incentive per se activations broadly agreed with results from the first GLM. RT effects (Figure 3D) then demonstrated associations with positive reward valence (both standard and jackpot trials), but not with negative valence (robber trials). On standard trials, we observed RT effects in multiple OLC sites (NBM, SA, and PUTv; Figure 3D), but not in amygdala. Among the CLC sites, only PUTd showed RT effects on these trials (Figure 3D). Additionally, RT effects on standard incentive trials were observed in a subset of our target motor areas (M1_UL_, M1_exUL_, SMA and CMA; Figure 3D) and in NAc. On jackpot trials, we observed RT effects in broadly similar sites, with two striking differences; we now observed RT effects in an amygdaloid OLC site (BLA), and we no longer observed the effect in any CLC sites (Figure 3D). On robber trials, in contrast, we observed no RT effects in any of our target sites (whether in OLC, CLC or a motor area). Interestingly however, robber trials were the only contexts associated with an RT effect in the opposite direction to all described above, that is, greater activation on trials with slower movement initiation. We observed a credible reverse RT effect solely on robber trials in a single motor area (PMv). In sum, by parsing RT effects across contexts, we observed that a subset of brain regions scaled activations with faster initiation time. However, involved subcortical regions were predominantly in the OLC and were only present on trials of positive valence (standard or jackpot). In contrast, we observed opposing RT effects in a separate motor site, on trials with negative valence incentives. Together these results implicate sites of OLC and a number of motor areas in potentially controlling the expedience of movement initiation when incentivized by pursuing reward but not avoiding loss.

## Discussion

As much as mammals rely on precision motor control to achieve goals in a complex world, the fundamental act of initiating a movement arguably remains the most important function of the motor system. Especially in light of the often transient windows of opportunity to capture reward or evade capture, it stands to reason that control circuitry linking motor areas in cortex to the striatum might have an in-built redundancy that augments control under special circumstances. In the present study we used an incentivized reaching task to explore whether one such redundancy might be an “open-loop” circuit involving sites of limbic and arousal function in addition to PUTv.

We first observed behaviorally that humans initiated precision reaches toward spatial targets more quickly if they were incentivized by high positive valence, as opposed to high negative valence. This faster movement initiation was accompanied by a greater likelihood of false starts, suggesting that high positive valence drove expedient responses over broader cognitive optimization. Alongside behavior, we observed an interesting interaction, whereby brain regions activated more for positive incentive cues at instruction phase of trials, then changed valence, activating more for negative incentive cues at the initiation of movement (go cue). This apparent temporal shift in neural responses to valence was observed in multiple regions including nodes of CLC, OLC and motor areas. We designed our models in such a way that these activations were separable neural correlates of behavior—specifically the expediency of movement initiation (RT). Recent simulations suggest that our approach can help avoid mistakenly concluding that different task conditions activate different brain regions, when in fact, the differences might be due to changes in reaction time (9). We exploited the reverse of this logic to reveal robust behavioral correlates—regions that scaled activation intensity with initiation time, or, “RT effects”. When collapsing across contexts, these RT effects were also in multiple regions of CLC, OLC and motor areas.

When we fitted a separate model that parsed behavioral correlates by context we revealed an important functional dissociation between OLC and CLC. Specific to high positive reward valence, RT effects were largely associated only with nodes in putative OLC alongside multiple regions of motor cortex. Beginning with the involved subcortical sites, we observed RT effects in BLA, consistent with the hypothesized inputs of the putative OLC, that is, projections from sites relevant for limbic function (15). In humans, the amygdala—and in particular, the BLA—have been shown to encode not just emotional valence but also the conjunction of emotional valence and action (16, 17). Critically, there is less degeneration of the BLA than central amygdala in patients with PD, underscoring a putative role for the BLA in generating paradoxical movement in this population (18). Connectomic evidence additionally supports BLA potentially serving as a crucial node transmitting goal-directed messages to the striatum. Specifically, the ventral part of the putamen (PUTv) is associated with areas relevant to affective and motivational processes, while the dorsal part of the putamen (PUTd) is instead linked with motor sites caudally and sites with roles in cognitive and executive functions rostrally (19, 20). We additionally observed RT effects in PUTv. Similar to BLA, PUTv has been shown to encode conjunctions of valence and action, with an emphasis on positive incentive and go behavior (21), similar to the context eliciting our observed effects. PUTv additionally appears to play a general role in goal-directed behavior, from learning to anticipate from previous experience (22, 23) to scaling the parameters of movement (24). Thus, although our observed effects in PUTv and BLA do not conclusively establish nodes of a neural circuit, they are consistent with literature suggesting that these regions are functionally connected and jointly implicated in relevant task settings.

In motor areas we observed the above RT effects (i.e., in contexts of high positive valence incentive) in M1, CMA and SMA. These motor areas are primarily involved in the planning and execution of movement, in addition to incorporating emotion and motivation with motor control. While we again cannot directly support a coordinated circuit with our data, effects in these motor areas are consistent with the hypothesized structure and function of OLC. First, anatomically, CMA and SMA in particular lie on medial walls of the frontal lobe, and appear to be functionally connected with putamen (20). Interestingly, more ventral divisions of putamen appear most connected with Brodmann area 26 (20), i.e., more proximal to CMA. This region, inferior to SMA, is thought to coordinate inner drive and affective responses into action in primates (25). This is supported by fMRI evidence in humans, showing that the CMA is more responsive to self-initiated and faster movements (26), which is closely related to our task. SMA itself is also associated with mechanisms relevant for our task, such as the timing of initiated movement (27, 28).

We here comment on two final observations. The first is the RT effects observed in NAc and NBM. NAc is strongly associated with reward anticipation in the incentive delay paradigm in humans (29, 30), and might even causally modulate motor areas (in particular SMA; 31). Here, we separated the classically defined NAc from other regions of the ventral striatum (PUTv) that appear to send prominent OLC projections to the motor cortical areas (3–6), as the function of these striatal regions has been relatively unexplored. Future work is needed to determine the extent to which the NAc contributes to the OLC and the intermediary regions in the OLC. The NBM is a promising candidate region for mediating limbic striatal outputs to the motor cortical areas ((4); Figure 1). In rodents, the NBM has been shown to have multisynaptic projections from PUTv to motor cortex (32). There is also evidence for PUTv projections to the NBM in the primate (33). It’s important to stress that the goal of the current study was to functionally dissociate putative OLC nodes from those in the classic CLC of movement initiation, and not assert that only one circuit modulates movement initiation speed. Future work will be needed to dissociate different aspects of OLC function (e.g., functions of glutamatergic neurons projecting from BLA to PUTv) from modulatory effects on motor control by dopaminergic and cholinergic systems, either within OLC itself or in separate circuits (e.g., striato-striatal, baso-striatal, etc.)

Second, we comment on the absence of RT effects on high negative valence incentive (robber) trials, and even the “reverse” RT effects observed in PMv. These results may suggest that positive reward incentive is the key context for driving OLC control of movement initiation. Indeed, participants likely responded to these contexts in psychologically different ways, despite their economic equivalence. This is evidenced first by the faster initiation of movement on jackpot trials, as well as the higher prevalence of false starts in that context. This is consistent with human learning and decision-making research, where findings frequently highlight an asymmetry between the influence of rewards and costs, both neurobiologically (34) and even in broader nervous system recruitment (35, 36). However, influential models of action selection also emphasize that it is unlikely a simple case that one circuit is activated by a context while another is not. Instead, multiple circuits likely compete for the ultimate control of movement (37). It’s entirely possible that OLC nodes might show RT effects to negative valence if there are no simultaneous activations of alternative circuits related to hesitation or stopping (38).

PD affects millions of people worldwide with no known cure. PK is observed in some individuals with PD under certain circumstances, but neither its substrate nor the exact contexts in which it occurs are fully understood. This study reveals task-based evidence that supports a putative OLC through basal ganglia, involving sites less impacted by the disease, that might selectively control movement under certain incentives. Future research involving PD patients will be necessary to verify if this circuit can facilitate movement in specific conditions when CLC is compromised. If successful, these findings could identify potential sites for therapeutic stimulation treatment.

## Acknowledgements

This research was funded in whole by Aligning Science Across Parkinson’s ASAP-020-519 through the Michael J. Fox Foundation for Parkinson’s Research (MJFF).

## Open Access Policy

All data used in the above analyses are freely available as part of the SKIP (SoCal Kinesia and Incentivization for Parkinson’s Disease) dataset through OpenNeuro.org and available under CC0 public copyright. For further information see www.socalkinesia.org. For the purpose of open access, the author has applied a CC0 public copyright license to all Author Accepted Manuscripts arising from this submission. Code for the incentivized reaching task can be obtained at https://doi.org/10.5281/zenodo.8216235. All ROI masks (Table 1) can be found in https://github.com/ejrise/asap_7T_resting.

**Table 1.**

| ROIs | Source |
| --- | --- |
| PMd, PMv, PUTd*, PUTv*, GPi, M1<sub>exUL</sub>**, SMA | Harvard Oxford atlas in (40) |
| NBM, SA | (42) |
| VL | (43) |
| M1<sub>UL</sub> | (44) |
| BLA, CeA | (45) |
| CMA | Hand-drawn by STG using (41) |

\*PUTd and PUTv obtained by segmenting putamen mask on the axial plane at $z = -1$ (MNI coordinates)
\*\*M1<sub>UL</sub> subtracted from this mask

## Materials and Methods

### Participants and testing procedure

We recruited 68 human participants (50 female; average age of 20.75 (SD=1.86)) via a combination of word-of-mouth and a study recruitment portal at UC Santa Barbara. All methods conducted were approved by the UC Santa Barbara Human Subjects Committee (protocol # 41-24-0102), which serves as the institutional review board for reviewing research applications involving human subjects. All participants gave informed written consent prior to participating.

Each subject’s data were collected in a single session, lasting approximately two hours. fMRI data were recorded by a Siemens 3T Prisma scanner with a 64-channel phased-array head/neck coil. Prior to entering the MR environment, participants completed a screening questionnaire to ensure they could enter safely. Task responses were recorded using an MR-compatible joystick. This was fixed to the center of a wooden table, which was placed across participants’ torsos once they were supine in the scanner bore. The position of the table was adjusted along the bore to ensure a comfortable distance for the participant’s arm. Arms were supported by pillows and padding.

Each session began with a a T1-weighted magnetization prepared rapid gradient echo (MPRAGE) anatomical scan (TR = 2500 ms, TE = 2.22 ms, FOV = 241 mm, T1 = 851 ms, flip angle = 7°, 0.94 mm^3^ voxel size). We next acquired a double-echo gradient echo field map sequence for distortion correction (TR = 758 ms, TE = 4.92 ms, flip angle = 60°, 2.5mm^2^ thick axial slices). We then acquired 4-dimensional echo-planar imaging during each of the three runs of the task (TR = 1900 ms, TE = 30 ms, flip angle = 65°, 2.5 mm^3^ voxel size, multiband acceleration factor of 2).

Task is described in the results section and in figure 1B. The deadline for reach completion from onset of the go cue (1.87 s) was calibrated from a pilot sample to achieve a 50% success rate across participants. This pilot sample of eight participants performed the task inside the scanner environment. From their data we used Bayesian estimation of the group-level mean and standard deviation of log-transformed completion times to infer the time associated with a 50% probability of completion.

Visual stimuli were rear-projected onto a screen placed approximately 110 cm behind participants using an LCD projector (1920×1080, 60 Hz). Participants viewed rectified images by way of a double mirror in the head coil. Each trial began with participants holding a screen cursor (radius=0.5 °; yoked to joystick position) at a starting position (radius=1.2 °; ∼9.5 ° below screen center). An instruction cue then appeared at one of two spatial locations. Each location had a radius of 1.2 ° and were both ∼9.5 ° above the screen center. The left target was ∼19 ° to the left of the screen’s midline, and the right target ∼19 ° to the right. Each instruction cue was made of the same small blue squares, re-arranged to make a neutral (standard trial), happy (jackpot) or sad (robber). Each stimulus was ∼6.6 ° radius and appeared centered on one of the two spatial targets for 200 ms. The hold period prior to the instruction cue (either 1 or 2 s) and go cue (1 - 4 s) altered across trials according to an m sequence. Total completion time on each trial was the time from the onset of the go cue until they reached the cued target, holding it in place for an additional 0.8 s. Completions within the deadline (1.87 s) were successful, with verbal feedback indicating the trial’s reward alongside the word “success”. Completions slower than this deadline were unsuccessful, with verbal feedback indicating the trial’s loss alongside the word “too slow”. All text was ∼2 ° in height and appeared at the screen center. Participants performed a set of 40 training trials prior to the experimental runs, where difficulty gradually increased from a deadline of 5 s to the task target.

### Behavioral measures

Cursor position was continuously recorded on each frame. Movement initiation time (RT) was defined as the time taken to initiate movement (depart the starting position) relative to the onset of the go-cue. False-starts were any movement outside of the starting position after the onset of the instruction cue, prior to the onset of the go cue. Behavior analysis of median RT, median completion time, average false-alarm rate and average trial success conducted with repeated measures ANOVA using factors of context (standard, jackpot, robber), hold period (>3 s,<=3 s) and run (1,2,3). Post-hoc pairwise correction for multiple comparison used FDR.

### fMRI preprocessing

We conducted MRI preprocessing with a custom pipeline featuring Advanced Normalization Tools (39) and FSL (40). T1-weighted anatomical data were skull-stripped with antsBrainExtraction.sh and resampled to functional image resolution (2 mm voxels). We then applied motion, slicetime, and distortion correction with field maps to EPI data with FSL FEAT. Next, we used ANTs to coregister and apply a non linear transform of EPI data to respective T1 anatomical data, then to MNI152 2mm space. EPI data were smoothed with a 5 mm FWHM Gaussian kernel.

*fMRI modeling* was performed using a combination of FSL and custom-made python scripts. Level 1 GLMs (i.e., run level) were fitted to each voxel’s preprocessed EPI time series using FSL’s feat. We fitted two GLMs presented in Figure 2A. The first GLM contained 7 regressors of interest. 3 stick function regressors (unit height, 0.1 s width) were event locked to the onset of the instruction cue, separate for each context. 3 stick function regressors (unit height, 0.1 s width) were event locked to the onset of the go cue, separate for each context. This allowed us to compute the four contrasts related to incentive per se reported in figure 3A-B (jackpot_inst_>standard_inst_; robber_inst_>standard_inst_; jackpot_go_>standard_go_; robber_go_>standard_go_). A duration scaled regressor (by RT) with unit height was event locked to the onset of the go cue. The contrast for RT effects was this coefficient compared to baseline (i.e., 1 vs 0s for all other regressors) which is reported in Figure 3C (RT_go_>baseline), i.e., activations that scaled intensity with faster RT. Note that this is a negative correlation, which we invert in reported results. A second GLM also had 7 regressors of interest. 3 stick function regressors (unit height, 0.1 s width) were event locked to the onset of the instruction cue, i.e., same as those used for the jackpot_inst_>standard_inst_ and robber_inst_>standard_inst_ contrasts in the first GLM. Three duration scaled regressors (by RT) with unit height were event locked to the onset of the go cue, separate for each context. The contrast for RT effects for each context were these coefficients compared to baseline (i.e., 1 vs 0s for all other regressors) which is reported in Figure 3D (standard_RTgo_>baseline; jackpot_RTgo_>baseline; robber_RTgo_>baseline). Each GLM contained two additional regressors of non interest. The first was unit height and duration scaled, with an onset at the offset of the instruction cue, and a duration until the onset of the go cue, i.e., the pre-go hold period. The second was also unit height and duration scaled, with an onset of the RT on each trial, and a duration until the end of the trial, i.e., reach activity. A level 2 GLM was then fitted to average the run-by-run activity in each voxel for each subject.

We hypothesized that specific regions are involved in the CLC and OLC, and sought to compare these two classes of regions. For a level 3 (i.e., across subjects) analysis, we therefore did not use a standard data-driven clustering approach of activations, and instead examined whether mean voxel activations in each ROI were modulated by the different task factors (contrasts). To this end, we used a hierarchical Bayesian model for each GLM. We first computed the average activation (Level 2) across voxels for each subject, for each of the ROIs described in Table 1, and for each contrast. We then assumed that the distribution of these averaged activations for each ROI-contrast pair across subjects (Level 3) could be described by a Student’s T distribution, where parameter nu would help accommodate outliers and provide more robust estimates of the central tendency (mu). For GLM1 a single mixture model fitted a mu, sigma and nu parameter for each of a total of 70 distributions (14 ROIs, 5 contrasts). Each distribution was assigned the same uninformed priors (mu_ROI,contrast_∼N(0,1); sigma_ROI,contrast_∼HalfN(1)). A single nu parameter accounted for all distributions and was also assigned an uninformed prior (nu∼HalfN(1)). For GLM2 a single mixture model fitted a total of 84 distributions (14 ROIs, 6 contrasts). We sought strong evidence for any central tendency parameter being credibly nonzero (p(mu_ROI,contrast_|y)>0) by requiring its entire highest density interval to be above or below 0. These values are reported in Figure 3.

## Glossary

BLA: basolateral amygdala
CeA: central nucleus of the amygdala
CLC: closed-loop circuit
CMA: cingulate motor area
GPi: globus pallidus internus
fMRI: functional magnetic resonance
M1_UL_: primary motor cortex, upper-limb region
M1_exUL_: primary motor cortex, excluding upper-limb region
NAc: nucleus accumbens
NBM: nucleus basalis of Meynert
OLC: open-loop circuit (putative)
PD: Parkinson’s disease
PK: paradoxical kinesia
PMd: dorsal premotor cortex
PMv: ventral premotor cortex
PUTd: dorsal (sensorimotor) putamen
PUTv: ventral (limbic) putamen
SA: septal area of the basal forebrain
SMA: supplementary motor area
VL: ventrolateral thalamus

## Notes

### Competing Interest Statement

The authors have declared no competing interest.

